# Epigenetically silencing gluconeogenic enzyme PCK1 by EZH2 promotes renal tubulointerstitial fibrosis

**DOI:** 10.1101/2022.10.31.514497

**Authors:** Ming Wu, Dongping Chen, Yanzhe Wang, Pinglan Lin, Feng Yang, Junyan Lin, Bo Tan, Guangyu Sheng, Xuejun Yang, Di Huang, Chaoyang Ye

## Abstract

Renal fibrosis is the common pathological pathway of various chronic kidney diseases (CKD) in progression to the end stage of renal failure. Impaired renal gluconeogenesis has been recently identified as a hallmark of CKD. The role of gluconeogenesis in renal fibrosis is currently unknown. Through RNA sequencing and cleavage under targets and tagmentation (CUT&Tag) sequencing analysis, we found that the phosphoenolpyruvate carboxykinase 1 (PCK1), a critical enzyme in gluconeogenesis, is negatively regulated by the methyltransferase enhancer of zeste homolog 2 (EZH2) in fibrotic kidneys, which was further confirmed by quantitative PCR, CUT&Tag and Western blotting analysis in unilateral ureteral obstruction (UUO) kidneys and renal cells. We further showed that pharmaceutical inhibition or conditional knockout of EZH2 increased PCK1 expression and attenuated renal fibrosis in two other mouse models. Moreover, the concentration of lactate, a glucogenic metabolic intermediate, was increased in mouse fibrotic kidneys, which was reduced by EZH2 inhibition. We further showed that glucose production was impaired in folic acid induced nephrotic mice and restored by EZH2 inhibition as assessed by pyruvate tolerance test. Importantly, the direct anti-gluconeogenesis effect of EZH2 on renal proximal epithelial cells was shown in human HK2 cells by analysis of gluconeogenic gene expression and production of glucose or lactate. Finally, inhibition of PCK1 by 3-mercaptopropionic acid abrogated the anti-fibrotic effect of EZH2 inhibitor in UUO kidneys. We conclude that EZH2 promotes renal tubulointerstitial fibrosis through epigenetic inhibition of PCK1. Our study suggests that therapeutic restoration of the impaired renal gluconeogenesis is beneficial to CKD patients.

**Translational Statement:** Renal fibrosis is the common pathological pathway of various chronic kidney diseases (CKD) in progression to the end stage of renal failure. Impaired renal gluconeogenesis has been recently identified as a hallmark of CKD. In this study, we found that the methyltransferase EZH2 is negatively correlated with the expression of the gluconeogenic enzyme PCK1 in fibrotic kidneys. Epigenetic inhibition of PCK1 by EZH2 promotes renal tubulointerstitial fibrosis. Our study indicates that renal gluconeogenesis in CKD can be epigenetically regulated and therapeutically targeted. Epigenetic restoration of renal gluconeogenesis is a potential therapeutic strategy to treat CKD patients with tubulointerstitial fibrosis.

## Introduction

Chronic kidney disease (CKD) affects more than 10% of global population, which is caused by acute or chronic injuries to glomerulus or renal tubules ^1,2^. The proximal tubule is the most vulnerable target of renal injuries and tubulointerstitial fibrosis is the best predictor of renal survival ^1,2^. Renal tubulointerstitial fibrosis is characterized by excessive deposition of extracellular matrix proteins (e.g. collagen-I and fibronectin), overexpression of epithelial-mesenchymal transition (EMT) markers (e.g. aSMA and N-cadherin), activation of pro-fibrotic signaling pathways (e.g. TGFβ/pSmad3 pathway) ^2,3^. Animal models have been established to study the pathogenesis of renal tubulointerstitial fibrosis by mimicking ischemic, obstructive, or toxic injuries in humans ^4,5^.

Epigenetic gene regulation by posttranslational modification of histone is involved in the development and disease condition of different organs ^6^. Enhancer of zeste homologue 2 (EZH2) is an important epigenetic regulator, and inhibition of EZH2 exerts renal protection in mouse models of acute kidney disease, lupus nephritis, and hyperuricemic nephropathy ^6^. EZH2 is up-regulated in unilateral ureter obstruction (UUO) kidneys and inhibition of EZH2 by 3-DZNeP attenuates renal tubulointerstitial fibrosis ^7,8^. However, the mechanism underlying the action of EZH2 in renal tubulointerstitial fibrosis is not completely known.

Gluconeogenesis is metabolic process by which glucose is synthesized from non-hexose precursors such as pyruvate and lactate ^9,10^. The gluconeogenesis pathway consists of 11 enzymes among which phosphoenolpyruvate carboxykinase (PCK1), fructose-1, fructose-1,6-bisphosphatase 1 (FBP1) and glucose-6-phosphatases (G6PC) were regarded as rate-limiting enzymes ^9^. Kidney is one of the major organs responsible for gluconeogenesis during fasting ^9^. Gluconeogenesis and glycolysis are two opposite metabolic pathways ^9^. Enhanced aerobic glycolysis in CKD is well-described and inhibition of glycolysis alleviates renal fibrosis in different animal models implying that a metabolic switch to gluconeogenesis might be beneficial to fibrotic kidneys ^9^. A very recent study showed that loss of renal gluconeogenesis is a hallmark of chronic kidney disease ^11^. Decreased renal gluconeogenesis was not only shown in fibrotic mouse kidneys but also associated with worse renal prognosis in CKD patients highlighting the importance of renal gluconeogenesis in the pathogenesis of chronic diseased kidneys ^11^. Whether improving renal gluconeogenesis can ameliorate renal tubulointerstitial fibrosis is currently unknown.

In this study, we found that renal gluconeogenesis is epigenetically controlled by EZH2 in fibrotic kidneys, which suggests that epigenetic restoration of renal gluconeogenesis is beneficial to CKD patients.

## Methods

### Animal models

Wide type male C57 mice were purchased from Shanghai Model Organisms Center Inc, and *Ezh2^fl/fl^* mice (referred to *Ezh2* mice) on a C57/B6 background were provided by Prof. Xi Wang from Capital Medical University ^12^. *Ezh2^fl/fl^* mice were crossed with C57/B6 tamoxifen-Cre (B6. Cg-Tg [Cre/Esr1]) 5Amc/J mice (stock 004682; Jackson Laboratories) as described previously ^13^. Animal experiments were approved by the ethic committee of Shanghai University of Traditional Chinese Medicine (PZSHUTCM18102604).

Unilateral ureteral obstruction (UUO) was established in mice (18-22g) as described before ^14^. *Ezh2* gene inactivation was induced by intraperitoneally injection of tamoxifen for three consecutive days at one week before sham or UUO operation. In brief, UUO operation was performed through twice ligation of the left ureter with 4-0 nylon sutures. WT mice (n= 7-8 mice per group) were treated with vehicle (normal saline) or 1 mg/kg 3-DZNeP (Apexibio, A8182) daily by i.p. injection. In some experiments, mice were treated with 30 mg/kg 3-Mercaptopropionic acid (M5801, Sigma-Aldrich) by gavage in combination with 1 mg/kg 3-DZNeP. Animals were sacrificed at day 7 or day 14 according to the experimental design.

For the unilateral ischemia-reperfusion injury (UIRI) model, left renal pedicles were clamped for 35 minutes by using microaneurysm clamps in male mice (18-22g). WT mice were randomly divided into three groups (n= 6-7 mice per group) and were treated with vehicle or 1 mg/kg 3-DZNeP daily by i.p. injection starting from day 1 for 10 days. Mice were sacrificed at day 11.

To establish folic acid (FA)-induced nephropathy, mice (25-30g, n= 6-10 mice per group) were injected intraperitoneally with FA (F7876, Sigma-Aldrich, dissolved in 0.3M sodium bicarbonate at a dose of 250 mg/kg). Sodium bicarbonate alone was used as control. *Ezh2* gene was inactivated at one week before FA injection. WT mice were treated with vehicle or 1 mg/kg 3-DZNeP daily by i.p. injection for 7 days starting from the operation day (day 0). At day 7 after FA administration, animals were sacrificed.

### Cell culture

Human kidney 2 (HK2) cells, a human renal proximal tubular epithelial cell line, were obtained from the Cell Bank of Shanghai Institute of Biological Sciences. HK2 cells were seeded in 6-well plate to 40-50% confluence, which were starved with DMEM/F12 medium 0.5% fetal bovine serum overnight before the experiment. On the next day, fresh medium containing 0.5% fetal bovine serum was changed, and then cells were exposed to 2.5 ng/ml TGF-β (Peprotech, Rocky Hill, NJ, USA) in the presence of DMSO or 20 μM of 3-DZNeP for a time duration according to the experimental design.

### RNA-seq Analysis

Mouse kidneys were harvested to prepare cDNA library, which was constructed by Biomarker Cloud Technologies and sequenced with paired-end reads on a Novaseq PE150 system. Detailed information was provided in the supplementary methods.

### Cleavage under targets and tagmentation (CUT&Tag) sequencing

Cells were collected to generate CUT&Tag libraries by Biomarker Technologies.

Primary antibody or IgG control was added to the cells bounded with balanced concanavalin A-coated magnetic beads. The Hyperactive Tn5 transposon fused with Protein A/Protein G was precisely targeted to cut the DNA sequence near the target protein. After DNA extraction, PCR amplification and PCR product purification, the library directly used for high throughput sequencing on an Illumina NovaSeq 6000 sequencer (Biomarker Technologies). Detailed information was provided in the supplementary methods.

### Quantitative PCR (qPCR)

Total RNA was extracted using Trizol (R401-01, Vazyme, Nanjing, China) from kidney samples according to the manufacture’s instruction, which was reverse transcribed to cDNA by Takara PrimeScript RT reagent kit (RR0036A, Kyoto, Japan). The primer sequences for qPCR were listed in Table S1. Hieff Unicon® qPCR TaqMan Probe Master Mix (11205E, Yeasen Biotechnology, Shanghai, China) was used for qPCR, which was performed using a StepOne Plus Sequence Detection System (Applied Biosystems).

### CUT&Tag-PCR Analysis

CUT&Tag experiments were performed following the procedures described in the CUT&Tag-seq section by using Hyperactive In-Situ ChIP Library Prep Kit for Illumina kit (Vazyme Biotech, TD901). Rabbit anti-EZH2 antibody (CST, 5246), the negative control rabbit IgG (CST, 3900) and the positive control Pol II antibody (CST, 2629) were used. Genomic DNA was purified by extraction with phenol:chloroform and amplified using specific primers (Table S2), which was further visualized on 2% agarose gels.

### Masson’s trichrome

Masson’s trichrome staining was performed using a standard protocol. Briefly, the four-μm-thick sections of paraffin-embedded kidney tissue was stained with hematoxylin, and then with ponceau red liquid dye acid complex, which was followed by incubation with phosphomolybdic acid solution. Finally, the tissue was stained with aniline blue liquid and acetic acid. Images were obtained with the use of a microscope (Nikon 80i, Tokyo, Japan).

### Western blotting analysis

Protein was extracted from mouse kidneys or HK2 cells, and were dissolved in 5x SDS-PAGE loading buffer (P0015L, Beyotime Biotech). After electrophoresis, proteins were electro-transferred to a polyvinylidene difluoride membrane (Merck Millipore, Darmstadt, Germany), which was incubated with first and secondary antibodies. Detailed information was provided in the supplementary methods.

### Gluconeogenic intermediate measurement

Renal cortex was collected for protein extraction. Protein samples were extracted by using 10 times normal saline through mechanical disruption. The content of lactate was measured with the lactate test kit (A019-2-1, NanJing JianCheng Bioengineering Institute, China) by determining OD value at 530 nm. The concentration of renal glucose was determined by measuring the OD value at 505 nm using glucose assay kit (F006-1-1) from NanJing JianCheng Bioengineering Institute. All measurements were performed according to the manufacturer’s instructions and were standardized by protein concentration.

HK2 human renal epithelial cells were starved overnight and followed by stimulation with TGF-β and treatment with 20 μM of 3-DZNeP. At 24h, supernatant was collected and centrifuged to remove the cell debris. The concentration of glucose or lactate was determined by commercial kits as described above.

### Pyruvate tolerance test

Mice were fasted for 16h before the test. Blood glucose levels were measured using a Roche Accu-Chek Active Blood Glucose Meter (Accu-Chek Active GB). Blood was sampled from the tail. Sodium pyruvate (A100342, Shanghai Shenggong Technology) was diluted in PBS, then 2g/kg sodium pyruvate solution was injected intraperitoneally. Glucose levels were recorded before injection and after 10, 20, 30, 60, 90 and 120min.

### Statistical analysis

Results were presented as mean ± SD. Differences among multiple groups were analyzed by one-way analysis of variance (ANOVA) and comparison between two groups was performed by unpaired student t-test by using GraphPad Prism version 8.0.0 for Windows (GraphPad Software, San Diego, California USA). A P value of lower than 0.05 was considered statistically significant.

## Results

### Deletion or inhibition of EZH2 attenuates renal tubulointerstitial fibrosis in obstructed mouse kidneys

We first studied the role of EZH2 in renal tubulointerstitial fibrosis by using *Ezh2* conditional knockout mice. No extracellular matrix deposition was observed in *Ezh2^fl/fl^;Cre/Esr1^-^* (control) or *Ezh2^fl/fl^;Cre/Esr1^+^* (*Ezh2*) conditional knockout mice with sham operation as shown by Masson staining (Figure 1A-1B). Masson staining showed that there was a strong renal interstitial fibrosis in control unilateral ureteral obstruction (UUO) mice, which was decreased in *Ezh2* knockout UUO kidneys (Figure 1A-1B). The protein levels of pro-fibrotic markers such as collagen-I, fibronectin, □SMA, and phosphrylation of Smad3 (pSmad3) were assessed by Western blotting analysis (Figure 1C-1H). UUO operation increased the expression of collagen-I, fibronectin, aSMA, and pSmad3 in control kidneys, which were decreased in *Ezh2* knockout UUO kidneys (Figure 1C-1H). The expression of renal EZH2 was also increased in control UUO kidneys, which were reduced in *Ezh2* knockout UUO kidneys (Figure 1C and 1F).

**Fig. 1.**
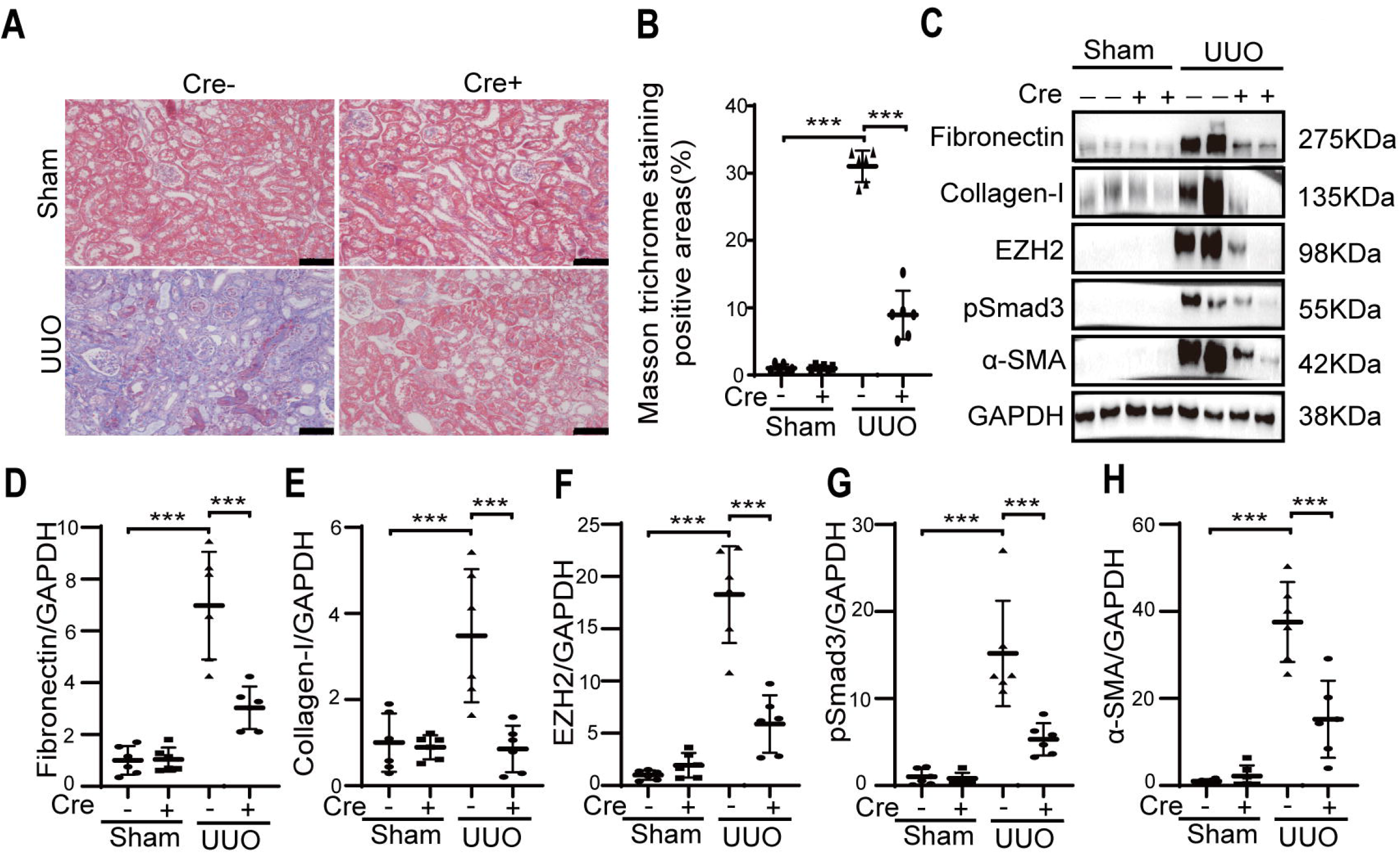
Deletion of EZH2 attenuates renal tubulointerstitial fibrosis in obstructed mouse kidneys. Control (*Ezh2^fl/fl^;Cre/Esr1^-^*) or (*Ezh2^fl/fl^;Cre/Esr1^+^*) *Ezh2* conditional knockout mice received sham or UUO operation and were sacrificed at day 14. (A-B) Renal fibrosis was assessed by Masson’s trichrome staining and further quantified. Scale bar represents 100 μm. (C-H) The expression of fibronectin, collagen-I, phosphorylated Smad3 (pSmad3), a-SMA and EZH2 were analyzed by Western blotting and further quantified. One representative of at least three independent experiments is shown. Data represent mean ± SD. \*\*\**p*□ <□0.001.

### EZH2 inhibits the expression of gluconeogenesis-related genes in UUO kidneys

To examine the effect of EZH2 activation on transcriptional output in fibrotic kidneys, we performed transcriptomic analyses by RNA sequencing (RNA-seq) in control and *Ezh2* knockout UUO kidneys. Total 768 differential expressed genes were identified by RNA-seq. Transcriptomic profiles of *Ezh2* knockout UUO kidneys differed substantially from those of control UUO kidneys, with 391 genes downregulated and 377 genes upregulated over 2-fold (Log2FC≥1) in *Ezh2* knockout UUO kidneys compared to control UUO kidneys (Figure 2A). To narrow downstream target genes of EZH2 in fibrotic kidneys, we performed Cleavage Under Targets and Tagmentation sequencing (CUT&Tag-Seq) in TGFβ treated human renal epithelial cells, which was combined with mRNA expression profiling (only 520 genes were mouse genes homologous to human). Through CUT&Tag-Seq, we found 15177 EZH2-binding genes. EZH2 is a well-known transcriptional repressor, thus we focused on 184 of the 222 upregulated genes in *Ezh2* knockout UUO group which had EZH2 occupancy (Figure 2B). KEGG Pathway analysis revealed a remarkable enrichment for genes linked to PI3K-Akt (*Sgk2; Gys2; Prlr; Fgf1*; *Pck1; G6pc*), PPAR (*Pck1; Slc27a2; Acox2*), Glucagon (*Gys2*; *Pck1; G6pc; Fbp1*), and AMPK (*Gys2*; *Pck1; G6pc; Fbp1*) signaling cascades (Figure 2C). We next examined the expression of several EZH2-target genes (*Sgk2; Gys2; Fgf1; Pck1; G6pc; Fbp1*) in UUO mouse kidneys from control and *Ezh2* knockout mice by performing quantitative PCR (qPCR) assays (Figure 2D). As shown in Figure 2E-2J, the expression of these EZH2-target genes in UUO kidneys were upregulated in *Ezh2* knockout mice. We next performed CUT&Tag-PCR with a focus on two rate-limiting enzymes of gluconeogenesis (PCK1 and G6PC). The binding of EZH2 to the promoter of these two genes was verified by CUT&Tag-PCR as compared with the negative IgG control and the positive control (RNA polymerase II RPB1) (Figure 2K). Western blotting analysis was performed to assess the expression of PCK1 and G6PC in the mouse UUO model. As compared with sham kidneys, the expression of PCK1 or G6PC was downregulated in UUO kidneys which was near or lower than the detection limit of Western blotting analysis (Figure 2L-2S). Western blotting analysis revealed the upregulation of PCK1 but not G6PC in UUO kidneys after genetic deletion of EZH2 (Figure 2L-2O). To further confirmed the inhibitory effect of EZH2 on PCK1 expression, we treated sham or UUO mice with the EZH2 inhibitor 3-DZNeP (Figure 2L-2S, and Figure S1. A-H). 3-DZNeP increased the expression of PCK1 but not G6PC in UUO kidneys, which was correlated with retarded progression of renal fibrosis in UUO mice as shown by Masson staining and Western blot analysis of collagen-I, fibronectin, aSMA, and pSmad3 expression (Figure S1. A-H).

**Fig. 2.**
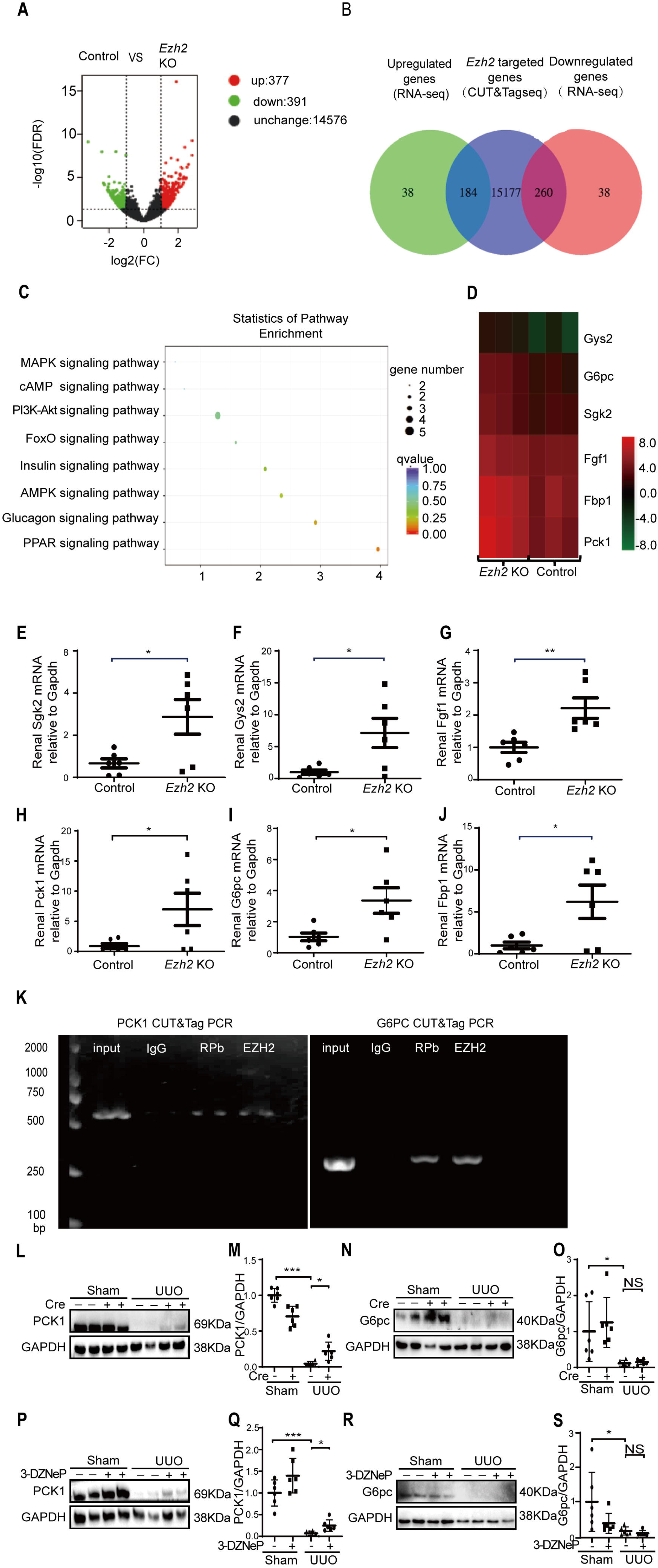
EZH2 inhibits gluconeogenic rate-limiting enzyme PCK1 in obstructed kidneys. (A) Volcano plots showing differentially expressed genes between control (*Ezh2^fl/fl^; Cre/Esr1^-^*) and (*Ezh2^fl/fl^; Cre/Esr1^+^) Ezh2* conditional knockout mice upon UUO operation, which was identified by RNA sequencing (RNA-seq). (B) Venn diagram showing the overlap of EZH2 regulated genes identified by RNA-seq and EZH2-binding genes which was identified by CUT&Tag sequencing in TGFβ stimulated human kidney epithelial (HK2) cells. (C) KEGG pathway enrichment analyses of EZH2-target genes. (D) Heatmap of expression values of genes enriched in several top regulated pathways by EZH2. Gene expression values are displayed by applying progressively brighter shades of red (up-regulated) or green (down-regulated). (E-J) Quantitative PCR analysis of mRNA levels of representative EZH2-target genes in kidneys between control and *Ezh2* conditional knockout mice upon UUO operation. (K) CUT&Tag PCR analysis of EZH2 occupancy on *PCK1* and *G6PC* genes in TGFβ stimulated HK2 cells. IgG was used as a negative control. Input and immunoprecipitates of anti-RNA polymerase II RPB antibody served as positive controls. (L-S) The expression of PCK1 and G6PC in EZH2 deleted or inhibited mouse sham or UUO kidneys were analyzed by Western blotting and quantified. These results are from three independent experiments. Data represent mean ± SD. NS represents not significant. \**p*□<□0.05. \*\**p*□<□0.01. \*\*\**p*□<□0.001.

### Ablation of EZH2 inhibits renal fibrosis and increases PCK1 expression in folic acid induced nephropathy

The role of EZH2 was further studied in a mouse model of folic acid (FA) nephropathy. Mouse FA nephropathy was induced by intraperitoneal injection of folic acid (250 mg/kg in 150 mmol/L sodium carbonate), and mice were sacrificed at day 7. As shown by Masson staining in Figure 3A-3B and 3J-3K, extracellular matrix was accumulated in FA mouse kidneys as compared with that in vehicle treated mouse kidneys, which was mitigated by genetic deletion or pharmaceutical inhibition of EZH2 in FA mice. Western blotting analysis revealed the upregulation of pro-fibrotic markers in FA kidneys as compared with that in vehicle treated WT or control kidneys, which was reduced by EZH2 inhibition or deletion (Figure 3C-3I and 3L-3R). Down-regulation of EZH2 and upregulation of PCK1 in FA kidneys by 3-DZNeP or genetic deletion of EZH2 were confirmed by Western blotting analysis (Figure 3C, 3F, 3L and 3O).

**Fig. 3.**
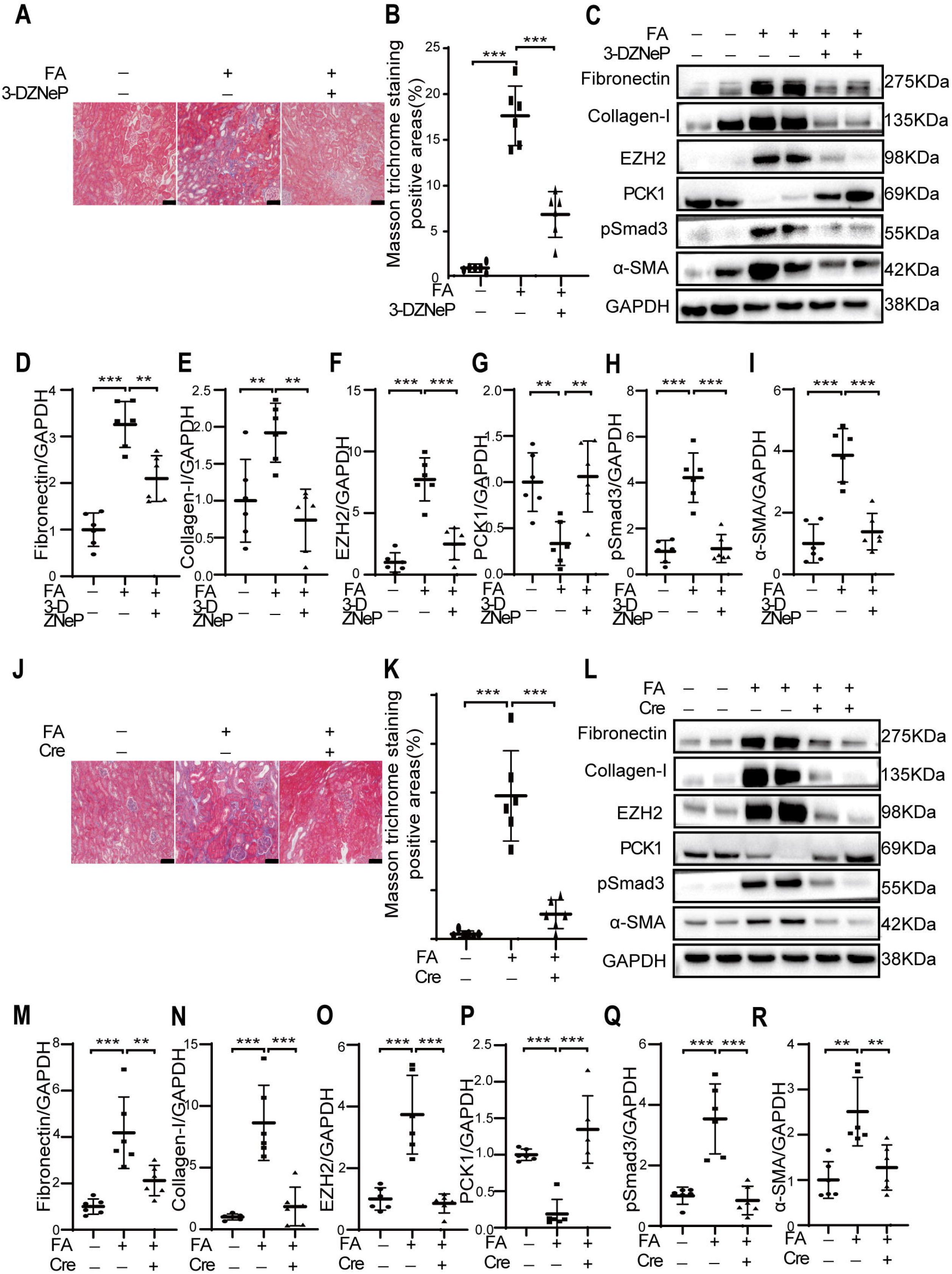
Ablation of EZH2 increases PCK1 expression and inhibits renal fibrosis in folic acid induced nephropathy. WT mice were injected with vehicle or folic acid (FA) intraperitoneally and treated with normal saline or 1 mg/kg 3-DZNeP for 7 days, which were sacrificed at day 7. (A-B) Renal fibrosis was assessed by Masson’s trichrome staining and further quantified. Scale bar represents 100 μm. (C-I) The expression of fibronectin, collagen-I, pSmad3, a-SMA, EZH2 and PCK1 were analyzed by Western blotting and further quantified. One representative of at least three independent experiments is shown; Control (*Ezh2^fl/fl^; Cre/Esr1^-^*) and (*Ezh2^fl/fl^;Cre/Esr1^+^) Ezh2* conditional knockout mice were injected with vehicle or FA intraperitoneally and were sacrificed at day 7. (J-K) Renal fibrosis was assessed by Masson’s trichrome staining and further quantified. Scale bar represents 100 μm. (L-R) The expression of fibronectin, collagen-I, pSmad3, a-SMA, EZH2 and PCK1 were analyzed by Western blotting and further quantified. One representative of at least three independent experiments is shown. Data represent mean ± SD. \*\**p*□ <□0.01. \*\*\**p*□ <□0.001.

### Ablation of EZH2 increases PCK1 expression and inhibits renal fibrosis in ischemia reperfusion induced nephropathy

The effect of EZH2 on PCK1 expression was further investigated by using the unilateral ischemia-reperfusion injury (UIRI) mouse model. The deposition of extracellular matrix proteins and the expression of pro-fibrotic markers were enhanced at day 11 after UIRI operation, which were attenuated by 3-DZNeP treatment as shown by Masson staining and Western blotting (Figure 4A-4I). In parallel, 3-DZNeP inhibited the expression of EZH2 and increased the expression of PCK1 in UIRI kidneys (Figure 4C and 4F).

**Fig. 4.**
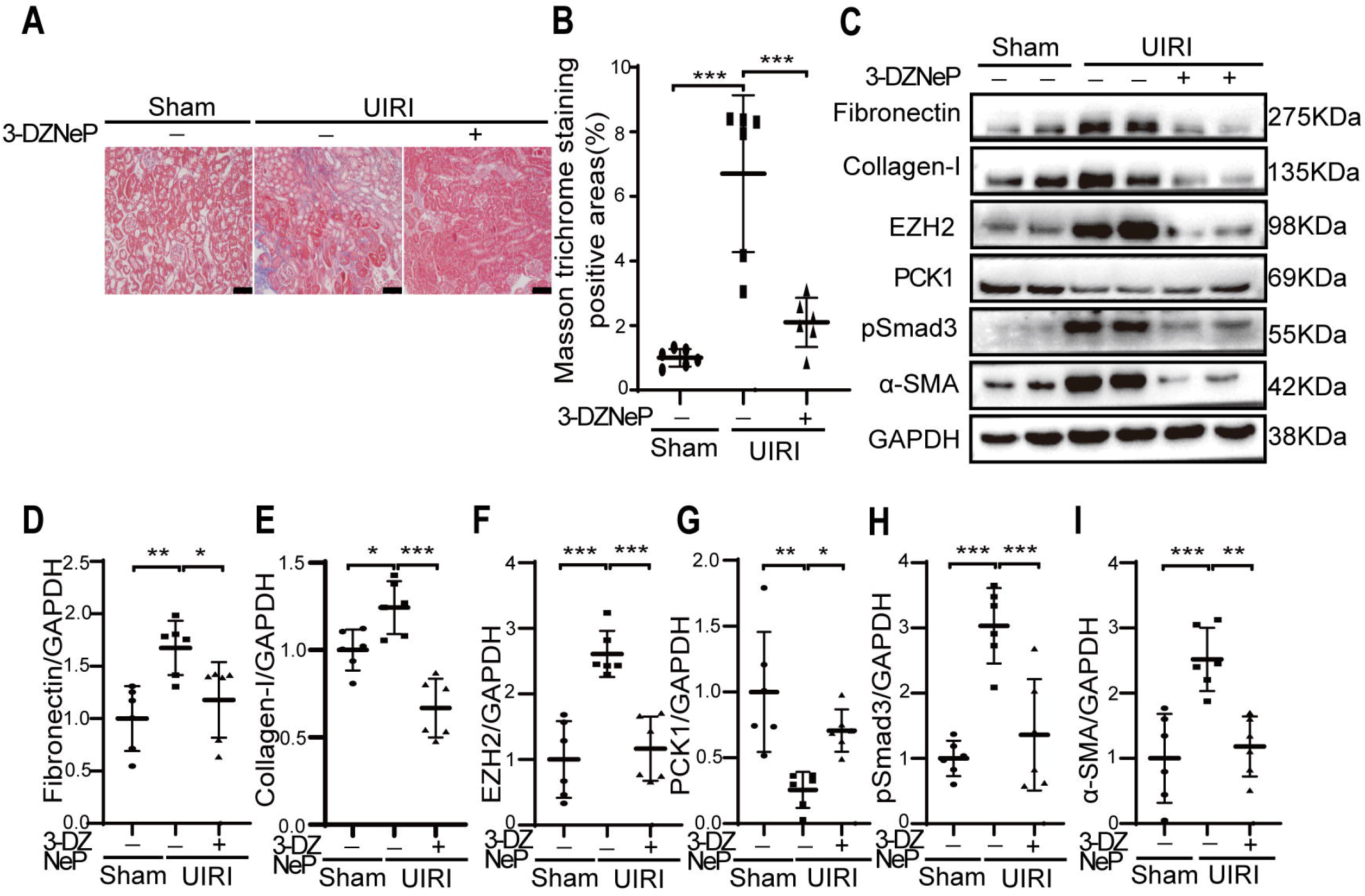
EZH2 inhibition increases PCK1 expression and inhibits renal fibrosis in ischemia reperfusion induced nephropathy. Wide type c57 mice underwent sham or unilateral ischemia-reperfusion injury (UIRI) operation, which were followed by treatment with normal saline or 1 mg/kg 3-DZNeP for 10 days starting from day 1. Mice were sacrificed at day 11. (A-B) Renal fibrosis was assessed by Masson’s trichrome staining and further quantified. Scale bar represents 100 μm. (C-I) The expression of fibronectin, collagen-I, pSmad3, a-SMA, EZH2 and PCK1 were analyzed by Western blotting and further quantified. One representative of at least three independent experiments is shown. Data represent mean ± SD. \**p*□ <□0.05. \*\**p*□ <□0.01. \*\*\**p*□ <□0.001.

### EZH2 inhibition improves renal gluconeogenesis in mice with fibrotic kidneys

Glucose metabolism were investigated in mice exposed to either sham or UUO operation at day 7. The difference in glucose concentration between sham and UUO kidneys was not observed (Fig. 5A). However, compared with sham mice, UUO mice had increased renal lactate concentration, and treatment with EZH2 inhibitor 3-DZNeP lowered lactate concentration in UUO kidneys (Fig. 5B). Renal glucose or lactate concentration was further measured in FA kidneys. However, the difference in glucose or lactate concentration between vehicle and FA kidneys was not observed (Figure S2 A-B). To further evaluation of glucose metabolism, we performed a pyruvate tolerance test in FA mice after injection of 2 g/kg sodium pyruvate. The increase in blood glucose levels following pyruvate injection was observed in control mice, which was peaked at 30 minutes (Figure 5C). Compared with control mice, the peak blood glucose level was significantly reduced in vehicle treated FA mice, which was returned to normal levels in 3-DZNeP treated FA mice (Figure 5C). Administration of FA reduced the area under the curve (AUC) and treatment of 3-DZNeP restored AUC to normal level (Figure 5D). All these data suggest that inhibition of EZH2 restores renal gluconeogenesis in mice with renal fibrosis.

**Fig. 5.**
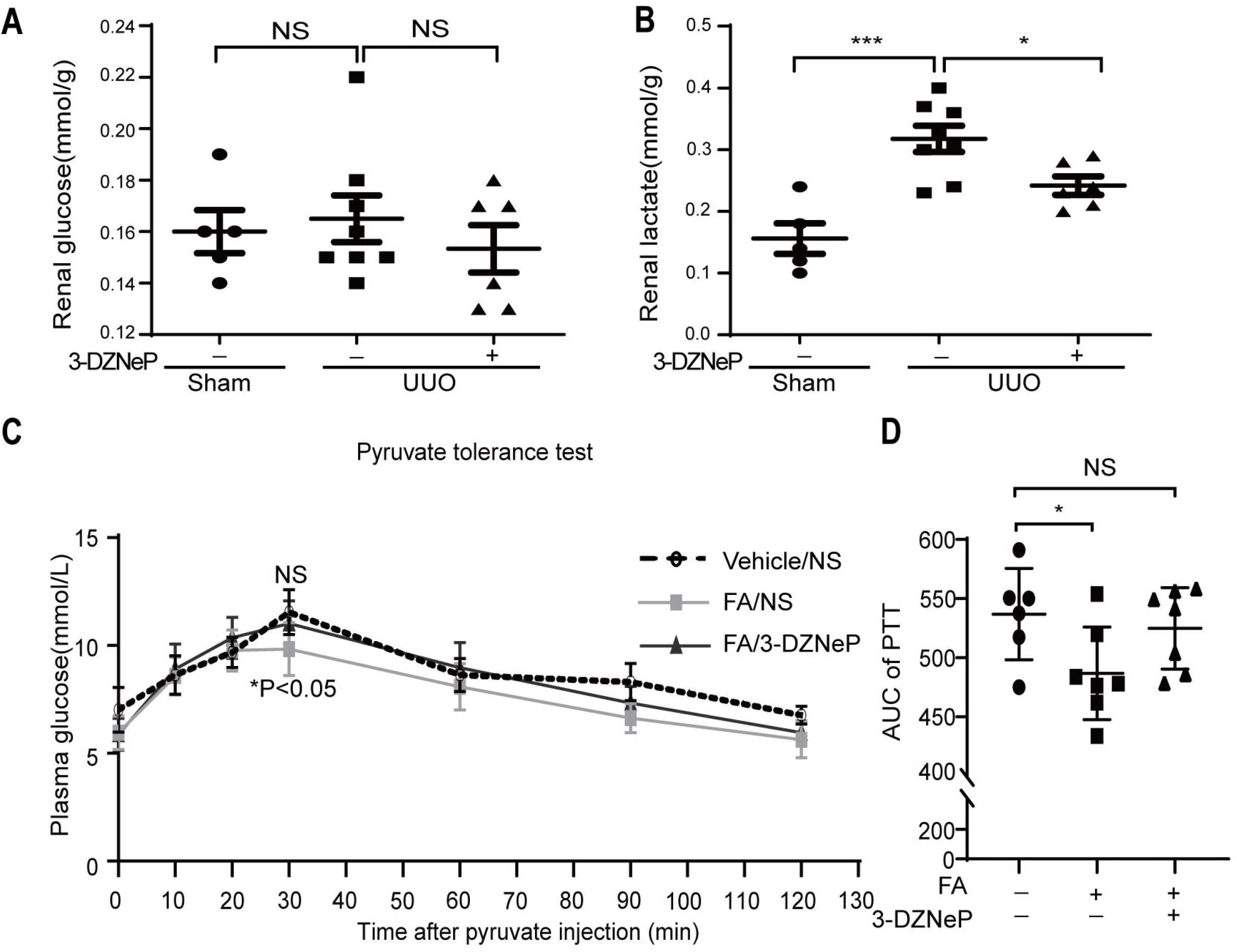
EZH2 inhibits gluconeogenesis in fibrotic kidneys. (A-B) WT mice received sham or UUO operation, which were followed by treatment with normal saline or 1 mg/kg 3-DZNeP for 7 days and were sacrificed at day 7. Mice were fasted for 16h before sacrifice. The levels of glucose and lactate in whole kidney samples were determined. Data are expressed as means ± SD (n = 5-8 mice per group). The statistical differences were evaluated using an unpaired t-test. NS represents not significant, \**p*□<□0.05. \*\*\**p*□<□0.001; WT mice received vehicle or folic acid (FA), which were followed by treatment with normal saline (NS) or 1 mg/kg 3-DZNeP for 7 days and were sacrificed at day 7. Mice were fasted for 16h before the pyruvate tolerance test (PTT). (C) Glucose levels in vehicle (n=□6 mice), FA (n=□7 mice) or FA with 3-DZNeP (n=□7 mice) were recorded at 0, 10, 20, 30, 60, 90, and 120 min after sodium pyruvate injection. (D) Area under the curve (AUC) of PTT. NS represents not significant, vehicle versus FA with 3-DZNeP at 30 min. \**p*□ <□0.05, vehicle versus FA at 30 min; One-way ANOVA was used.

### EZH2 inhibits gluconeogenesis of renal epithelial cells in a fibrotic condition

To examine the direct anti-gluconeogenic effect of EZH2 on renal epithelial cells in a fibrotic condition, we tested the effect of EZH2 inhibitor 3-DZNeP on TGFβ stimulated HK2 human renal proximal epithelial cells. Firstly, we investigated the time course expression of three gluconeogenic genes (PCK1, G6PC and FBP1) in TGFβ stimulated HK2 cells. As shown in Figure 6A-6C, 1h TGFβ stimulation reduced the mRNA expression of PCK1 and G6PC but not FBP1 in HK2 cells. Treatment with 3-DZNeP reversed the mRNA expression of PCK1 and G6PC in TGFβ stimulated HK2 cells at 1h (Figure 6D-6E). We next assessed the effect of 3-DZNeP on lactate and glucose metabolism in TGFβ stimulated HK2 cells. Compared with control cells, TGFβ stimulation increased lactate concentration and decreased glucose concentration in the supernatant of HK2 cells, which were reversed by 3-DZNeP treatment (Figure. 6F-6G). Inhibition of EZH2 expression and pro-fibrotic marker expression (fibronectin, N-cadherin, and pSmad3) by 3-DZNeP were confirmed in TGFβ stimulated HK2 cells by Western blotting analysis (Figure 6H-6L). Together, these data suggest that EZH2 inhibits gluconeogenesis of renal epithelial cells in a fibrotic condition.

**Fig. 6.**
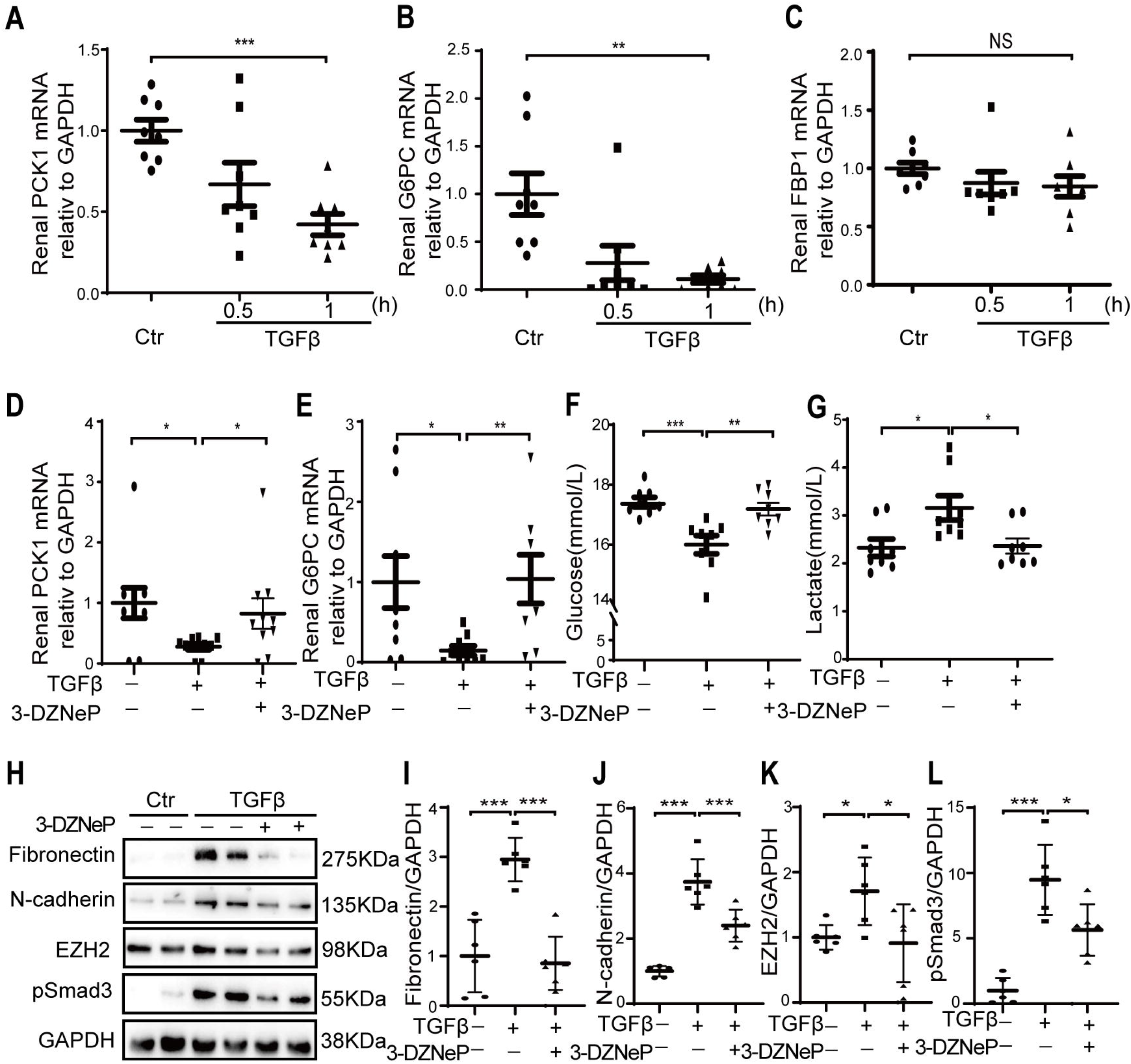
EZH2 inhibits gluconeogenesis of renal epithelial cells in a fibrotic condition. (A-C) HK2 human renal epithelial cells were starved overnight and followed by stimulation with TGF-β for 0.5 to 1h. The mRNA expression of PCK1, G6PC and FBP1 were analyzed by qPCR. (D-E) HK2 human renal epithelial cells were starved overnight and followed by stimulation with TGF-β and treatment with 20 μM of EZH2 inhibitor 3-DZNeP for 1 h. The mRNA expression of PCK1 and G6PC were analyzed by qPCR. (F-G) HK2 human renal epithelial cells were starved overnight and followed by stimulation with TGF-β and treatment with 20 μM of 3-DZNeP for 24h. The concentration of glucose and lactate in the supernatant collected and were determined. (H-L) HK2 human renal epithelial cells were starved overnight and followed by stimulation with TGF-β and treatment with 20 μM of 3-DZNeP for 48h. Cell lysates were extracted. The expression of fibronectin, N-cadherin, pSmad3 and EZH2 were analyzed by Western blotting and then quantified. Data represents mean□±□SD. One representative result of at least three independent experiments is shown. NS represents not significant. \**p*□<□0.05. *\*\*p*□<□0.01. \*\*\**p*□<□0.001.

### Inhibition of PCK1 prevents the anti-fibrotic effect of EZH2 inhibitor

PCK1 inhibitor 3-Mercaptopropionic acid (3MPA) was used in UUO models to test whether EZH2 promotes renal fibrosis through PCK1 ^15^. Renal fibrosis was assessed at one week after UUO operation by Masson staining or Western blotting. Reduced deposition of extracellular matrix proteins and decreased expression of pro-fibrotic markers can be observed in 3-DZNeP treated UUO kidneys, and the anti-fibrotic effect of 3-DZNeP in UUO kidneys was abolished by the treatment of 3MPA (Figure 7A-7G).

**Fig. 7.**
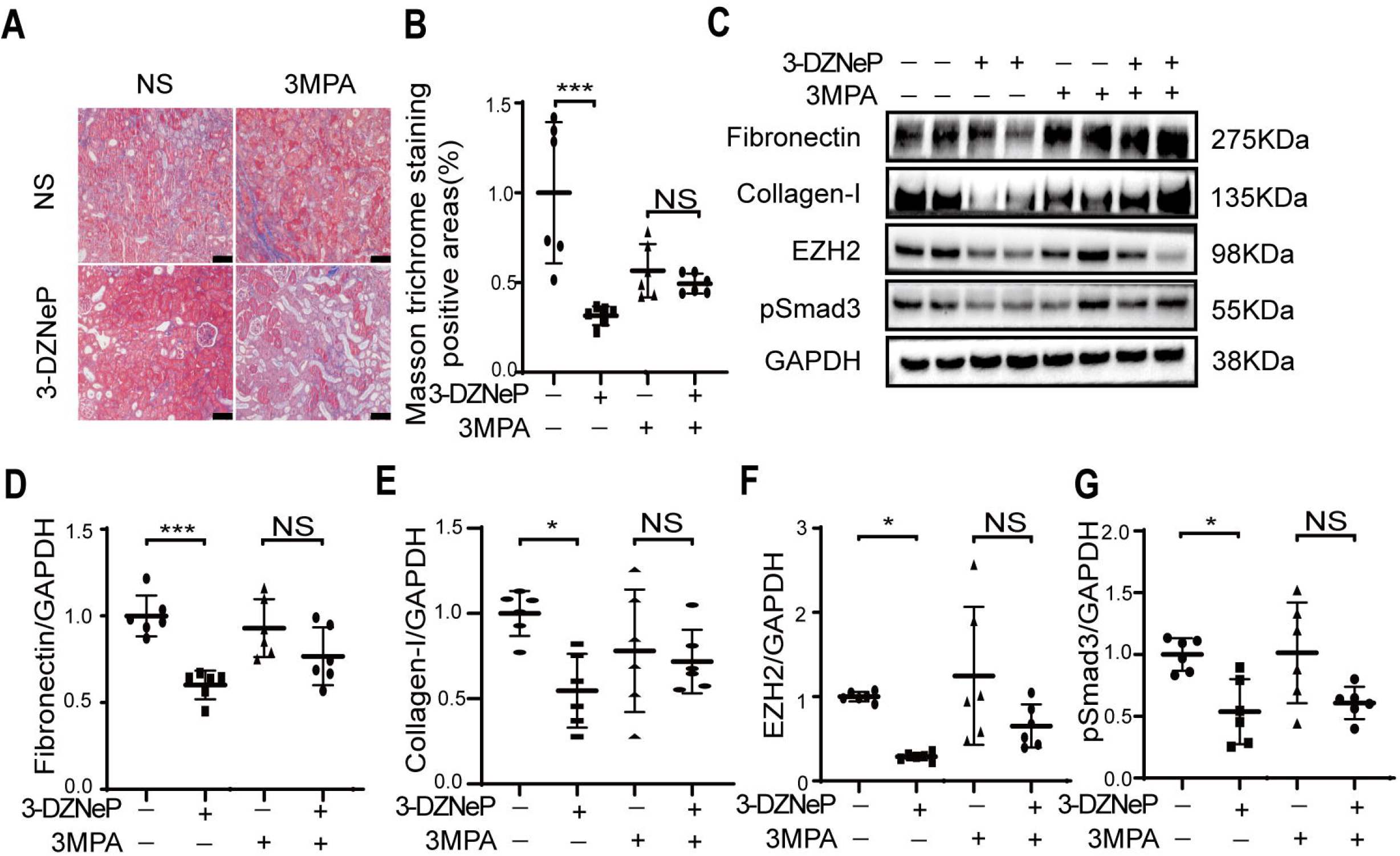
Inhibition of PCK1 reverses the anti-fibrotic effect of EZH2 inhibitor 3-DZNeP in fibrotic kidneys. WT mice received UUO operation, which were followed by treatment with normal saline (NS) or 1 mg/kg 3-DZNeP in combination with 30 mg/kg PCK1 inhibitor 3-Mercaptopropionic acid (3MPA) or NS for 7 days. Mice were sacrificed at day 7. (A-B) Renal fibrosis was assessed by Masson’s trichrome staining and then quantified. Scale bar represents 100 μm. (B) The expression of fibronectin, collagen-I, pSmad3, and EZH2 were analyzed by Western blotting and further quantified. One representative of at least three independent experiments is shown. Data represent mean ± SD. NS represents not significant. \**p*□<□0.05. \*\*\**p*□<□0.001.

## Discussion

Previous study showed that pharmaceutical inhibition of EZH2 mitigates renal tubulointerstitial fibrosis in UUO mice ^7,8^. In the current study, we confirmed the anti-fibrotic effect of EZH2 by using conditional knockout mice in UUO model. The conclusion was further proved by using pharmaceutical and genetic approach in ischemia reperfusion or nephrotoxin induced renal tubulointerstitial mouse models.

It has been shown that EZH2 inhibition attenuates renal tubulointerstitial fibrosis by activation of PTEN/Akt signaling pathway ^8^. To further understand the mechanism underlying the pro-fibrotic effect of EZH2 in renal tubulointerstitial fibrosis, we performed RNA-seq and CUT&Tag-Seq to search direct downstream targets of EZH2. We found that the expression of several gluconeogenesis-related genes is negatively regulated by EZH2 in UUO kidneys, which were further validated by qPCR in *Ezh2* conditional knockout UUO kidneys. By using CUT&Tag-PCR, we verified the binding of EZH2 on the promoter of two rate-limiting enzymes of gluconeogenesis (PCK1 and G6PC). We further showed the expression of PCK1 protein was downregulated in three mouse models of renal tubulointerstitial fibrosis and inhibition or deletion of EZH2 reversed PCK1 expression in fibrotic mouse kidneys. Moreover, inhibition of EZH2 by 3-DZNeP reduced the accumulation of lactate, the gluconeogenic intermediate, in UUO kidneys. Furthermore, EZH2 inhibition restored renal gluconeogenesis in FA mice as evaluated by pyruvate tolerance test. Finally, we showed that inhibition of PCK1 by 3MPA abolished the anti-fibrotic effect of 3-DZNeP in UUO kidneys. We conclude that EZH2 promotes renal tubulointerstitial fibrosis through inhibition of the key gluconeogenesis enzyme PCK1. Thus, in this study we revealed a novel mechanism underlying the anti-fibrotic effect of EZH2 in kidney disease.

More importantly, our study is the first study showing that restoration of renal gluconeogenesis is beneficial to chronic kidney disease. Impaired renal gluconeogenesis occurs early in CKD patients and becomes worse in parallel to the progression of CKD stages ^11^. Renal gluconeogenesis is negatively correlated with renal fibrosis in CKD patients and our study suggests that therapeutic targeting renal gluconeogenesis could be a potential strategy to inhibit renal fibrosis in CKD patients ^11^. Renal gluconeogenesis also has direct systemic effect on CKD patients. Impaired renal gluconeogenesis could induce hypoglycemia or hyperlactatemia, which are associated with high mortality and poor renal prognosis in CKD patients and ICU patients ^10,11^. Recovery of renal gluconeogenesis could improve the mortality rate of CKD patients through systemically restoring blood glucose and lactate levels.

Transcriptional regulation of gluconeogenesis is well studied and several transcriptional factors such as peroxisome proliferator-activated receptor alpha (PPARα) can regulate gluconeogenic gene expression ^9^. Epigenetic regulation of gluconeogenesis is a new research topic and has been studied in recent years. Repression of gluconeogenic gene expression by EZH2 has recently been shown in liver and kidney cancer ^16^. EZH2 is enriched in the promoter of FBP1 and promotes tumor growth through repression of FBP1 ^16^. Importantly, the inversed correlation of EZH2 and FBP1 in tumor samples can accurately predict patient survival ^16^. This is the first time, we showed that renal gluconeogenesis can be epigenetically regulated in kidney diseases, suggesting a novel intervention direction to improve renal gluconeogenesis. The protective effect of SGLT2 inhibitors or thiamine supplement shown by clinical studies in CKD or ICU patients may partly rely on their ability to improve renal gluconeogenesis ^10^. Whether EZH2 is involved in the beneficial effect of both treatments is not known. Several clinical trials using EZH2 inhibitors are undergoing in cancer patients and specifically targeting EZH2 mediated epigenetic regulation of renal gluconeogenesis in CKD patients is expecting.

Renal gluconeogenesis takes place in renal proximal tubules ^10^. One limitation in our study is that *Ezh2* gene is ubiquitously deleted in our conditional knockout mice by using the CAG-Cre/Esr1 system, which is a ubiquitous system and is not kidney specific ^17^. We are unable to conclude the direct anti-gluconeogenic effect of EZH2 on renal proximal epithelial cells based on our *in vivo* data. To answer this question, we established an *in vitro* model by using HK2 cells, a well-known human renal proximal tubular cell line, to study the effect of EZH2 on renal gluconeogenesis. Upon TGFβ stimulation, two gluconeogenic genes are downregulated and the EZH2 inhibitor reversed the expression of these two genes. Importantly, we observed that reduced glucose and increased lactate production upon TGFβ stimulation, which were reversed by EZH2 inhibition. More importantly, the improved renal gluconeogenesis *in vitro* was correlated with reduced expression of pro-fibrotic markers. Thus, we conclude that EZH2 directly inhibits gluconeogenesis of renal proximal tubular cells in a fibrotic condition, which may contribute to the anti-fibrotic effect of EZH2.

In conclusion, EZH2 promotes renal tubulointerstitial fibrosis through inhibition of the gluconeogenic enzyme PCK1 expression, suggesting that epigenetic regulation of gluconeogenesis could be a strategy to treat CKD patients.

## Supplementary Material

### Supplementary Methods

#### RNA-seq Analysis

Mouse kidneys were harvested from three control and three *Ezh2* knockout mice at day 14 after UUO operation. Total RNA was isolated from mouse kidneys using TRIzol (Life technologies, California, USA), and subsequently cDNA library was constructed by Biomarker Cloud Technologies and sequenced with paired-end reads on a Novaseq PE150 system. Low quality reads were removed from the raw RNA-seq reads using perl script. Clean reads were mapped with the mouse genome using Hisat2 (V2.2.1). Differentially expressed genes were calculated by DESeq2_EBSeq algorithms with FDR (False Discovery Rate) ≤ 0.05 and fold-change >2. Function enrichment analysis was performed using KOBAS 3.0. All raw data have been deposited under the Gene Expression Omnibus accession number SUB10878122.

#### Cleavage under targets and tagmentation (CUT&Tag) sequencing

HK2 cells were starved overnight and were exposed to 2.5 ng/ml TGFβ (Peprotech, Rocky Hill, NJ) for 48h. Cells were collected and frozen in media consisting of 10%DMSO and 20% FBS, which were sent to Biomarker Technologies to generate CUT&Tag libraries.

Balanced concanavalin A-coated magnetic beads bound to 60-100,000 prepared cells by incubating at room temperature. Rabbit anti-EZH2 antibody (CST, 5246) or rabbit IgG (CST, 3900) was then added to the beadbound cells. The Hyperactive Tn5 transposon fused with Protein A/Protein G was precisely targeted to cut the DNA sequence near the target protein through the mediation of the respective incubation of the primary antibody, the corresponding secondary antibody and Protein A/Protein G. DNA was sheared by the tagmentation buffer by incubation at 37□. After DNA extraction, PCR amplification and PCR product purification, the library directly used for high throughput sequencing on an Illumina NovaSeq 6000 sequencer (Biomarker Technologies). For CUT&Tag data processing, low-quality reads and adapters were filtered by the Cutadapt software. The clean reads were mapped to human genome with Bowtie2 software. MACS2 v2.1.1 software was further used to detect genome-wide Peak region. The ChIPseeker package was used for functional annotation of genome-wide Peak, and DiffBind package was used for Difference Peak analysis. All raw data have been deposited under the Gene Expression Omnibus accession number SUB10926540.

#### Western blotting analysis

Renal protein was extracted from mouse kidneys or HK2 cells by using RIPA lysis buffer (P0013B) from Beyotime Biotechnology (Nantong, China). The protein concentration was measured by the Bradford method, and the supernatant was dissolved in 5x SDS-PAGE loading buffer (P0015L, Beyotime Biotech). Samples were subjected to SDS-PAGE gels. After electrophoresis, proteins were electro-transferred to a polyvinylidene difluoride membrane (Merck Millipore, Darmstadt, Germany), which was incubated in the blocking buffer (5% non-fat milk, 20mM Tris-HCl, 150mMNaCl, PH=8.0, 0.01%Tween 20) for 1 hour at room temperature and was followed by incubation with anti-fibronectin (1:1000, ab23750, Abcam), anti-pSmad3 (1:1000, ET1609-41, HUABIO), anti-Collagen I (1:500, sc-293182, Santa Cruz), anti-α-SMA (1:1000, ET1607-53, HUABIO), anti-N-cadherin (1:1000, ET1607-53, HUABIO), anti-Snail (1:1000, ET1607-53, HUABIO), anti-GAPDH (1:5000, 60004-1-lg, Proteintech), or anti-α-tubulin (1:1000, AF0001, Byotime) antibodies overnight at 4□. Binding of the primary antibody was detected by an enhanced chemiluminescence method (BeyoECL Star, P0018A, Byotime) using horseradish peroxidase-conjugated secondary antibodies (goat anti-rabbit IgG, 1:1000, A0208, Beyotime or goat anti-mouse IgG, 1:1000, A0216, Beyotime). The quantification of protein expression was performed using -Image J.

**Fig. S1.** Inhibition of EZH2 attenuates renal tubulointerstitial fibrosis in obstructed mouse kidneys.

WT mice received sham or UUO operation, which were followed by treatment with normal saline (NS) or 1 mg/kg 3-DZNeP for 14 days and were sacrificed at day 14. (A-B) Renal fibrosis was assessed by Masson’s trichrome staining and further quantified. Scale bar represents 100 μm. (C-H) The expression of fibronectin, collagen-I, pSmad3, a-SMA and EZH2 were analyzed by Western blotting and further quantified. One representative of at least three independent experiments is shown. Data represent mean ± SD. \**p*□ <□0.05. \*\*\**p*□ <□0.001.

**Fig. S2.** Gluconeogenesis in folic acid induced fibrotic kidneys.

WT mice received vehicle or folic acid (FA) and were sacrificed at day 7. Mice were fasted for 16h before sacrifice. The levels of (A) glucose and (B) lactate in whole kidney samples were determined. Data are expressed as means ± SD (n = 8-9 mice per group). The statistical differences were evaluated using an unpaired t-test. NS represents not significant.

## References

1 Chevalier, R. L. The proximal tubule is the primary target of injury and progression of kidney disease: role of the glomerulotubular junction. Am J Physiol Renal Physiol 311, F145–161, doi:10.1152/ajprenal.00164.2016 (2016).

2 Gewin, L. S. Renal fibrosis: Primacy of the proximal tubule. Matrix Biol 68-69, 248–262, doi:10.1016/j.matbio.2018.02.006 (2018).

3 Schnaper, H. W. The Tubulointerstitial Pathophysiology of Progressive Kidney Disease. Adv Chronic Kidney Dis 24, 107–116, doi:10.1053/j.ackd.2016.11.011 (2017).

4 Fu, Y. et al. Rodent models of AKI-CKD transition. Am J Physiol Renal Physiol 315, F1098–F1106, doi:10.1152/ajprenal.00199.2018 (2018).

5 Chevalier, R. L., Forbes, M. S. & Thornhill, B. A. Ureteral obstruction as a model of renal interstitial fibrosis and obstructive nephropathy. Kidney Int 75, 1145–1152, doi:10.1038/ki.2009.86 (2009).

6 Li, T., Yu, C. & Zhuang, S. Histone Methyltransferase EZH2: A Potential Therapeutic Target for Kidney Diseases. Front Physiol 12, 640700, doi:10.3389/fphys.2021.640700 (2021).

7 Zhou, X. et al. Targeting histone methyltransferase enhancer of zeste homolog-2 inhibits renal epithelial-mesenchymal transition and attenuates renal fibrosis. FASEB J, fj201800237R, doi:10.1096/fj.201800237R (2018).

8 Zhou, X. et al. Enhancer of Zeste Homolog 2 Inhibition Attenuates Renal Fibrosis by Maintaining Smad7 and Phosphatase and Tensin Homolog Expression. J Am Soc Nephrol 27, 2092–2108, doi:10.1681/ASN.2015040457 (2016).

9 Faivre, A., Verissimo, T., Auwerx, H., Legouis, D. & de Seigneux, S. Tubular Cell Glucose Metabolism Shift During Acute and Chronic Injuries. Front Med (Lausanne) 8, 742072, doi:10.3389/fmed.2021.742072 (2021).

10 Legouis, D., Faivre, A., Cippa, P. E. & de Seigneux, S. Renal gluconeogenesis: an underestimated role of the kidney in systemic glucose metabolism. Nephrol Dial Transplant 37, 1417–1425, doi:10.1093/ndt/gfaa302 (2022).

11 Verissimo, T. et al. Decreased Renal Gluconeogenesis Is a Hallmark of Chronic Kidney Disease. J Am Soc Nephrol 33, 810–827, doi:10.1681/ASN.2021050680 (2022).

12 Yin, J. et al. Ezh2 regulates differentiation and function of natural killer cells through histone methyltransferase activity. Proceedings of the National Academy of Sciences of the United States of America 112, 15988–15993, doi:10.1073/pnas.1521740112 (2015).

13 Sun, Y. et al. Activation of P-TEFb by cAMP-PKA signaling in autosomal dominant polycystic kidney disease. Sci Adv 5, eaaw3593, doi:10.1126/sciadv.aaw3593 (2019).

14 Wu, M. et al. Reduced asymmetric dimethylarginine accumulation through inhibition of the type I protein arginine methyltransferases promotes renal fibrosis in obstructed kidneys. FASEB J 33, 6948–6956, doi:10.1096/fj.201802585RR (2019).

15 Ma, R. et al. A Pck1-directed glycogen metabolic program regulates formation and maintenance of memory CD8(+) T cells. Nat Cell Biol 20, 21–27, doi:10.1038/s41556-017-0002-2 (2018).

16 Liao, K. et al. A Feedback Circuitry between Polycomb Signaling and Fructose-1, 6-Bisphosphatase Enables Hepatic and Renal Tumorigenesis. Cancer Res 80, 675–688, doi:10.1158/0008-5472.CAN-19-2060 (2020).

17 Donocoff, R. S., Teteloshvili, N., Chung, H., Shoulson, R. & Creusot, R. J. Optimization of tamoxifen-induced Cre activity and its effect on immune cell populations. Sci Rep 10, 15244, doi:10.1038/s41598-020-72179-0 (2020).

