## Supplemental Table for "Epigenetically silencing gluconeogenic enzyme PCK1 by EZH2 promotes renal tubulointerstitial fibrosis"

**Table S1: List of primers used for quantitative PCR**

| Primer Name | Forward | Reverse |
|---|---|---|
| **mouse Sgk2** | GAGACCACGTCCACCTTCTG | CGTAAGGCTCTTTACGAAGCA |
| **mouse Gys2** | CAGCACCATTAGACGAATCGGAC | CCAAGGTGACAACCTCGGACAA |
| **mouse Fgf1** | GTAGTTTCCTAGAGGCAGGTTG | TGATAAAGTGGAGTGAAGAGAGC |
| **mouse Pck1** | CTCAGCTGGCAGCATGGGGTG | AACAGCTCCTCCACGTTGACG |
| **mouse G6pc** | CCACTTTGGTTTCAGTTGAATCAG | GTATCCACCAGTAAGGACGATGG |
| **mouse Fbp1** | GCT CTG CAC CGC GAT CA | ACA TTG GTT GAG CCA GCG ATA |
| **mouse Gapdh** | AGGTCGGTGTGAACGGATTTG | TGTAGACCATGTAGTTGAGGTCA |
| **human PCK1** | GGTTCCCAGGGTGCATGAAA | CACGTAGGGTGAATCCGTCAG |
| **human G6PC** | CCTCAGGAATGCCTTCTACG | TCTCCAATCACAGCTACCCA |
| **human FBP1** | GATTGCCTTGTGTCCGTTG | TGCCATACAGTGCGTAGCC |
| **human GAPDH** | GAGTCAACGGATTTGGTCGT | TTGATTTTGGAGGGATCTCG |

**Table S2: List of primers used for CUT&Tag-PCR**

| Primer Name | Forward | Reverse |
|---|---|---|
| **human PCK1** | AGTCTGAATGATTTGCTTCACAG | CCTCAACAGATCCACGTACCC |
| **human G6PC2** | ACTGATCTCAGGTGGAACACAG | CTACGACTGGACCCTGGACTA |
