## Supplementary figures and images for "Epigenetically silencing gluconeogenic enzyme PCK1 by EZH2 promotes renal tubulointerstitial fibrosis"

### Supplemental Figure1

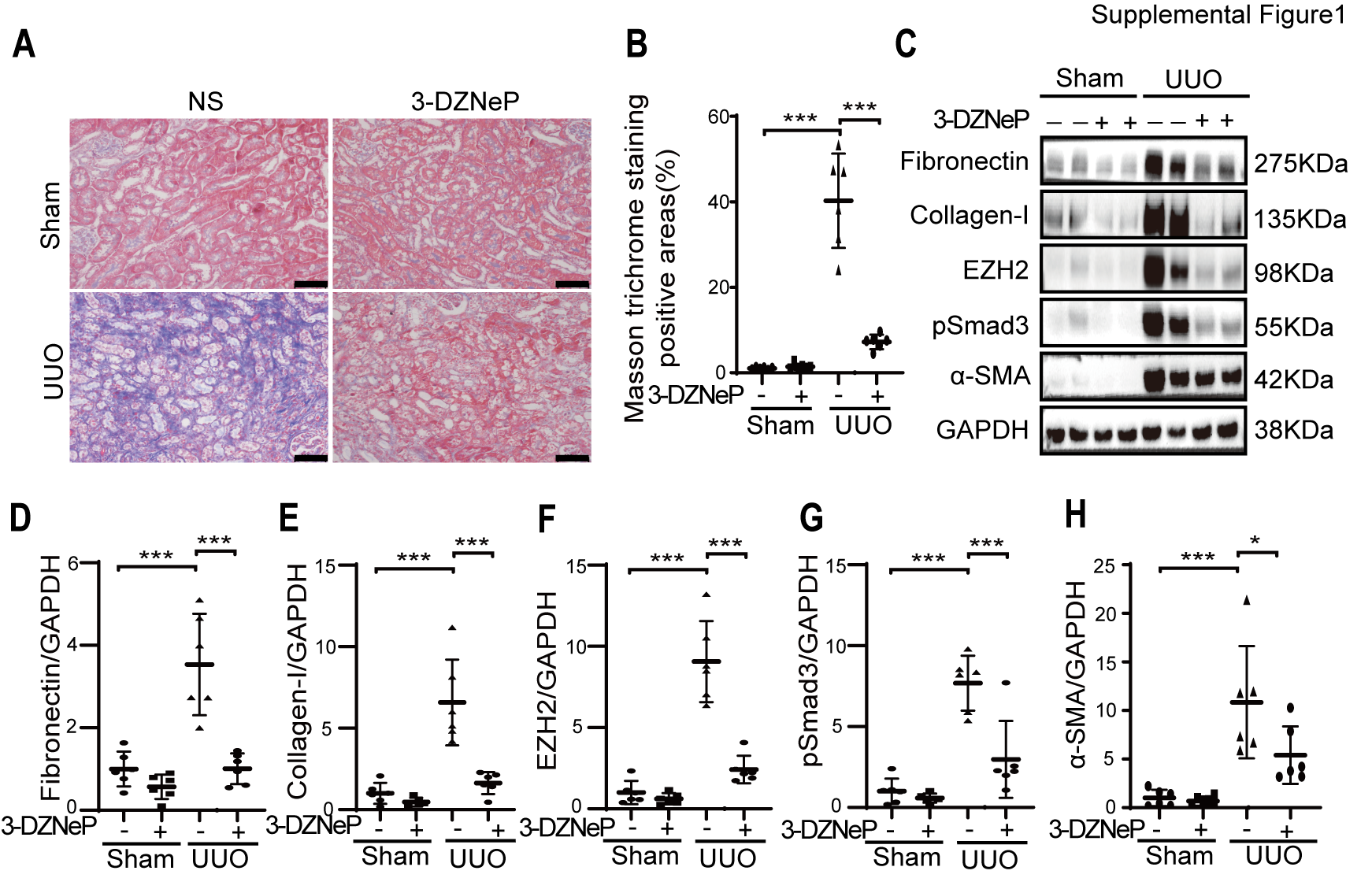

### Supplemental Figure2

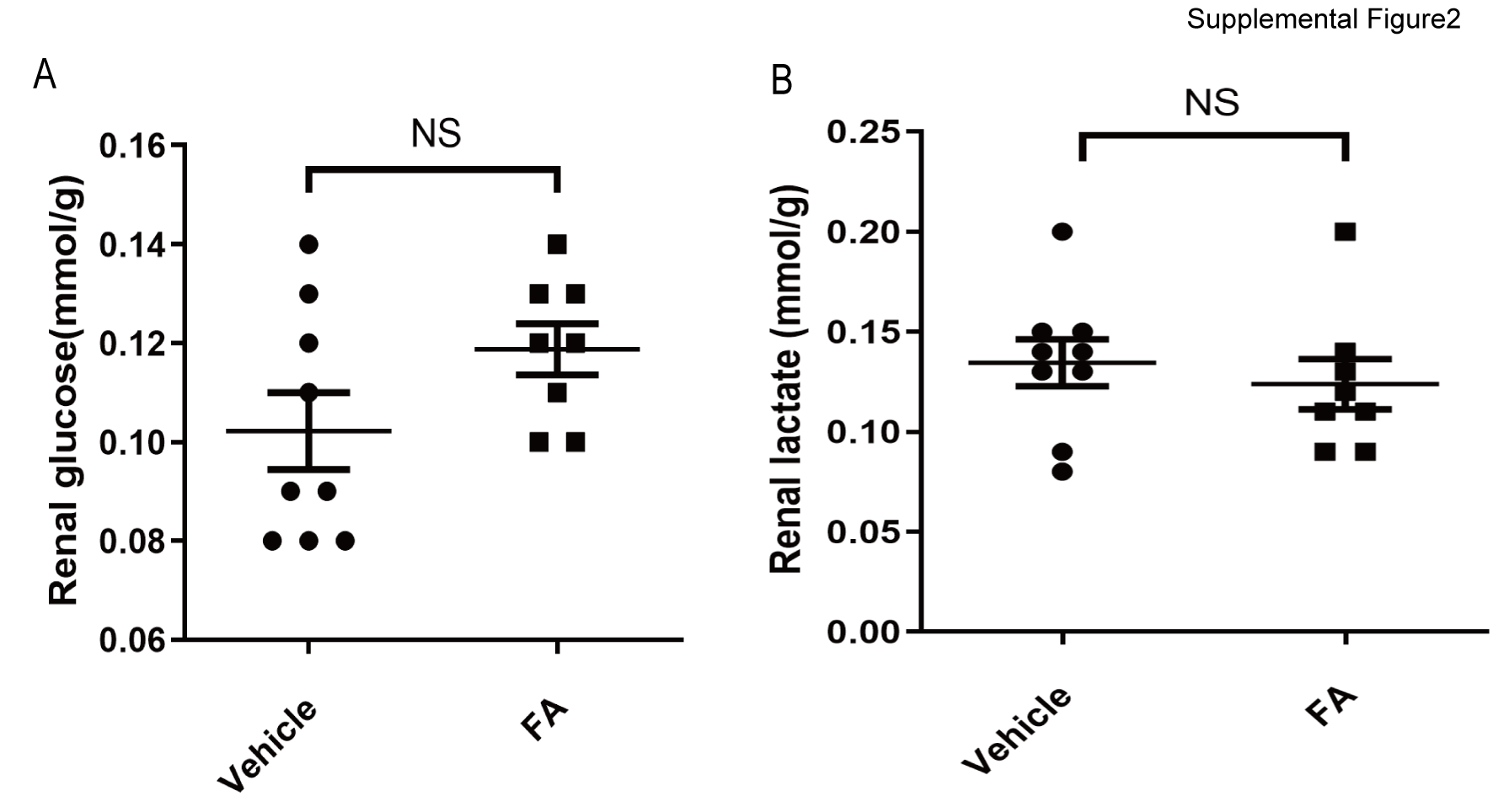
